# Non-spherical coacervate shapes in an enzyme driven active system

**DOI:** 10.1101/714261

**Authors:** Willem Kasper Spoelstra, Eli O. van der Sluis, Marileen Dogterom, Louis Reese

## Abstract

Coacervates are polymer-rich droplets that form through liquid-liquid phase separation in polymer solutions. Liquid-liquid phase separation and coacervation have recently been shown to play an important role in the organization of biological systems. Such systems are highly dynamic and under continuous influence of enzymatic and chemical processes. However, it is still unclear how enzymatic and chemical reactions affect the coacervation process. Here, we present and characterize a system of enzymatically active coacervates containing spermine, RNA, free nucleotides, and the template independent RNA (de)polymerase PNPase. We find that these RNA coacervates display transient non-spherical shapes, and we systematically study how PNPase concentration, UDP concentration and temperature affect coacervate morphology. Furthermore, we show that PNPase localizes predominantly into the coacervate phase and that its depolymerization activity in high-phosphate buffer causes coacervate degradation. Our observations of non-spherical coacervate shapes may have broader implications for the relationship between (bio-)chemical activity and coacervate biology.

## Introduction

Complex coacervation is the chemical process in which a solution of oppositely charged polymers phase separates into a coacervate phase (polymer-enriched) and a solvent phase (polymer-depleted). It is one of multiple forms of polymeric liquid-liquid phase separation (LLPS), which occurs when the interactions between polymers dominate over the entropy of mixing^1–4^. The recent discovery that membraneless organelles in cells exhibit liquid-like properties^5^ gives rise to a plethora of novel biological, chemical and biochemical questions^6–9^. Previous studies have also highlighted the importance of intracellular LLPS in (neurodegenerative) diseases, including Amyotrophic Lateral Sclerosis (ALS)^10,11^ and Alzheimer’s disease^12^. Understanding the roles of LLPS in biological systems now presents an emerging challenge in the life sciences.

It is in the nature of living systems to constantly proliferate, implying that their components are constantly replicated using internal and external energy sources. Chemical and enzymatic reactions facilitate the continually dynamic state of cells. Here we are interested in active coacervates, which are coacervates with external or internal energy supply (see Berry et al.^13^ and Weber et al.^14^ for recent reviews). Active coacervate systems have been studied by Alexander Oparin, who hypothesized that such systems provided a stepping stone for the origin of life on earth^15–20^. He provided insightful examples about model systems for membraneless biological compartmentalization. Oparin’s ideas have been revisited in experiments but faded into the background, potentially owing to the difficulty of characterizing coacervates^21^.

Due to the recent discovery of intracellular LLPS^5^, coacervates have regained attention in both experimental^22–29^ and theoretical^30–32^ work. For example, studies have shown that onset of coacervation can be reversibly regulated by enzymatically altering the phosphorylation state of the components (e.g. peptides^23^, ATP^26^). Additionally, it has been established that some coacervates facilitate RNA catalysis^27^ and can regulate template-directed RNA polymerization at sub-optimal magnesium levels^29^. Furthermore, theory has shown that active coacervate droplets can display shape instabilities under specific conditions, for example when droplet material is continuously being produced outside and degraded inside of the coacervate phase^31,32^. However, it has remained unclear how active and passive coacervates differ in terms of their physicochemical properties. Here, we introduce coacervate morphology as a property affected by enzyme activity. We use a model system where active RNA/spermine coacervates display non-spherical shapes, as opposed to passive spherical coacervates. The system is driven by the enzymatic polymerization reaction of Polynucleotide Phosphorylase (PNPase) on RNA templates. We observe that these active coacervates initially are non-spherical and we quantify the time for which the assumed non-equilibrium shapes persist, and assess their degree of non-sphericality. To compare contributions of active and passive processes in the system, we also quantify the timescale of merging for passive coacervates and the timescale of non-equilibrium shape deformation. We find that these timescales are separated by two orders of magnitude. The relaxation of non-equilibrium shapes occurs within several minutes, whereas merging of two coacervates is a matter of a few seconds. Moreover, we assessed the localization of the PNPase and found that the PNPase localizes predominantly, but not entirely, inside the coacervate phase. Taken together, this work provides characterization of a model system that can be exploited to model intracellular biomolecular condensation and test emerging theories of active LLPS.

## Materials & Methods

All experiments were carried out in 1x PNPase synthesis buffer (100 mM Tris-HCl, 1 mM EDTA, 5 mM MgCl_2_, pH 9.0)^33^, and nuclease free water was used in all experiments. Poly(U)_∞_ was synthesized by mixing 5 µM Cy5-poly(U)_20_, 40 mM UDP, and 4 µM PNPase in 1x PNPase synthesis buffer, and left at 30 °C for >2 hours. In contrast, we use “poly(U)” without the lemniscate subscript to refer to commercial polyuridylic acid which was of unspecified length. Polyvinyl alcohol, spermine tetrahydrochloride, Trizma hydrochloride, magnesium dichloride (MgCl_2_), sodium chloride (NaCl), potassium chloride (KCl), (Ethylenedinitrilo)tetraacetic acid (EDTA), uridine diphosphate disodium salt (UDP), uridine monophosphate disodium salt (UMP), polyuridylic acid potassium salt (poly(U)), poly(U)_20_ and Cy5-poly(U)_20_ were bought from Sigma-Aldrich. Nuclease free water, High Range Riboruler, SYBR Safe, fluorescein-5-maleimide were purchased from Thermo Fisher.

### Sample preparation

For imaging, glass coverslips were coated with polyvinyl alcohol (PVA). First, the coverslips were exposed to oxygen plasma (Plasma Preen, Plasmatic Systems) for 15 seconds, and subsequently they were submerged into a 5% (w/v) PVA solution and left for 5 minutes at room temperature. The PVA-coated coverslips were then dried under a flow of nitrogen and placed on top of channels cut into parafilm before heating to 120 °C for five minutes. Slides were used within 24 hours. Due to the hydrophilic nature of this PVA coating, the wetting of the glass surface by the coacervate phase was prevented (**Fig. S2**) and coacervates remained mobile (**Supporting Videos**).

### Confocal microscopy

Coacervate samples were imaged using a Nikon Eclipse Ti confocal microscope with perfect focus system, an oil immersion objective (Nikon Plan Apo λ 100x NA 1.45), an EMCCD camera (Photometrics Evolve 512), a spinning disc unit (CSU-W1, Yokogawa), and the FRAP/TIRF system Ilas2 (Roper Scientific) for illumination. A custom-made objective heater was used for temperature control of the samples. Unless stated otherwise, the temperature was fixed at 30 °C across all experiments.

### Image processing

Binarization of coacervate-micrographs for circularity analysis was done by smoothing and thresholding in ImageJ (default, v1.52g)^34^. Then the shape descriptors were calculated, from which the circularity of all detected objects were extracted and plotted to obtain Fig. 2d-f. Objects on the edges and objects smaller than 36 pixels (0.92 µm^2^ cross sections) were excluded from analysis. The number of coacervates analyzed and their size (by cross-sectional area) for each condition tested in Figures 2d-f, are shown in **Figure S3**.

### CRAM-analysis

For the CRAM-analysis (Fig. 3), merging events of coacervates were recorded at 272 ms intervals. These recordings were manually cropped such that only the two merging coacervates were fully inside the cropped region (**Fig. S4a**). Images were then smoothed and thresholding was applied (see **Image processing**) to obtain binary images of merging coacervates (**Fig. S4b**). For each image, the area and circularity of the coacervates was calculated. Time zero was defined as the frame from which only a single coacervate was detected. To quantify the circularity (*t* ≥ 0), we fitted the single exponential function

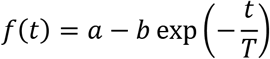

where *T*, *a* and *b* are free fitting parameters. *T* represents the characteristic timescale of merging, which was plotted against the radius of the final coacervate to obtain Fig. 3c. The slope of this data is the inverse capillary velocity, which we determined by fitting a straight line through the origin. The difference (*a* − *b*) represents the fitted circularity at *t* = 0, and the parameter *a* is the average circularity of the spherical coacervates. A geometric argument (**Fig. S4c**) yields that for the merging of two equal-sized and perfectly spherical coacervates, *a* = 1 and 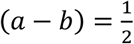. The obtained values and statistics of the fitting parameters are presented in (**Fig. S4d**).

### PNPase-localization and FRAP measurements

For the localization study of PNPase (Fig. 4), FITC-PNPase was spun down at 30 psi to remove aggregates. A mixture of 4 µM FITC-PNPase, 1.0 wt% spermine, 20 mM UDP and 5 µM Cy5-poly(U)_20_ was prepared in 1x PNPase synthesis buffer. Fluorescence micrographs were acquired using confocal microscopy and analyzed to obtain local fluorescence intensities and intensity ratios. To calculate the fluorescence intensities in both phases, the Cy5-poly(U) fluorescence intensities were used as a mask. Micrographs acquired by capturing Cy5-poly(U) fluorescence were binarized (see **Image Processing**) and these binary images were multiplied with the micrographs acquired by capturing FITC-PNPase fluorescence. From the images produced by this multiplication, we calculated the average fluorescence intensity, discarding black pixels. For detailed description of the workflow, see **Figure S6**. This exact same workflow was used to extract the background intensity of FITC fluorescence in both phases and at the interface, and these values were subtracted from the FITC-PNPase intensity calculated from the experiment to obtain Figure 4c. From these values we calculated the partition coefficient of FITC-PNPase into the coacervate phase using

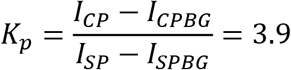

This partition coefficient was converted into a partitioning free energy via^35^

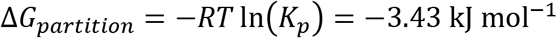

The Fluorescence Recovery After Photobleaching (FRAP) analysis was done by fully bleaching the FITC-PNPase inside a square around the spherical coacervate, using a 488 nm wavelength laser. The FITC-PNPase intensity was recorded and normalized by the average pre-bleach intensity to obtain Figure 4d.

## Results

### Enzymatic activation of RNA/spermine coacervates

It is well established that long poly(U) RNA and spermine undergo LLPS by forming complex coacervates^36^. Long homopolymeric poly(U) RNA can be formed by the enzyme Polynucleotide Phosphorylase (PNPase). In vivo, PNPase plays a role in RNA metabolism^37–42^. In vitro, PNPase adds uridine monophosphate nucleotides (UMP) to the 3’-end of RNA seeds when provided with uridine diphosphate (UDP) (Fig. 1a and **Fig. S1**). We first investigated whether active poly(U)/spermine coacervates form by enzymatic polymerization from short (20 nts) poly(U) seeds, as schematically shown in Figure 1b. We observed that adding PNPase to a mixture of poly(U)_20_, spermine, and UDP, coacervates formed within 15 minutes, as the solution turns turbid. If UDP was substituted by UMP, no turbidity was observed, confirming that coacervation was the result of PNPase-mediated poly(U) polymerization (Fig. 1c & d). To study the emergence of this enzymatically-triggered phase separation, we monitored the formation of poly(U)/spermine with confocal microscopy. Because PNPase binds and polymerizes RNA at the 3’-end, we used a fluorescent label (Cy5) at the 5’-end of the poly(U)_20_ seeds. The reaction mixtures were prepared and flushed into a passivated glass channel for imaging (**Materials & Methods**). Within five minutes, the coacervates sedimented onto the bottom of the channel. Notably, we observed non-spherical shapes in the first 10-15 minutes of the experiment instead of spherical droplets (Fig. 1e). Non-spherically shaped coacervates are only possible in a non-equilibrium situation. Here the directionality of RNA polymerization by PNPase appears sufficient to maintain non-equilibrium for a limited amount of time, while presence of the polyelectrolyte spermine condenses the polymerized poly(U) RNA into an active coacervate.

**Figure 1:**
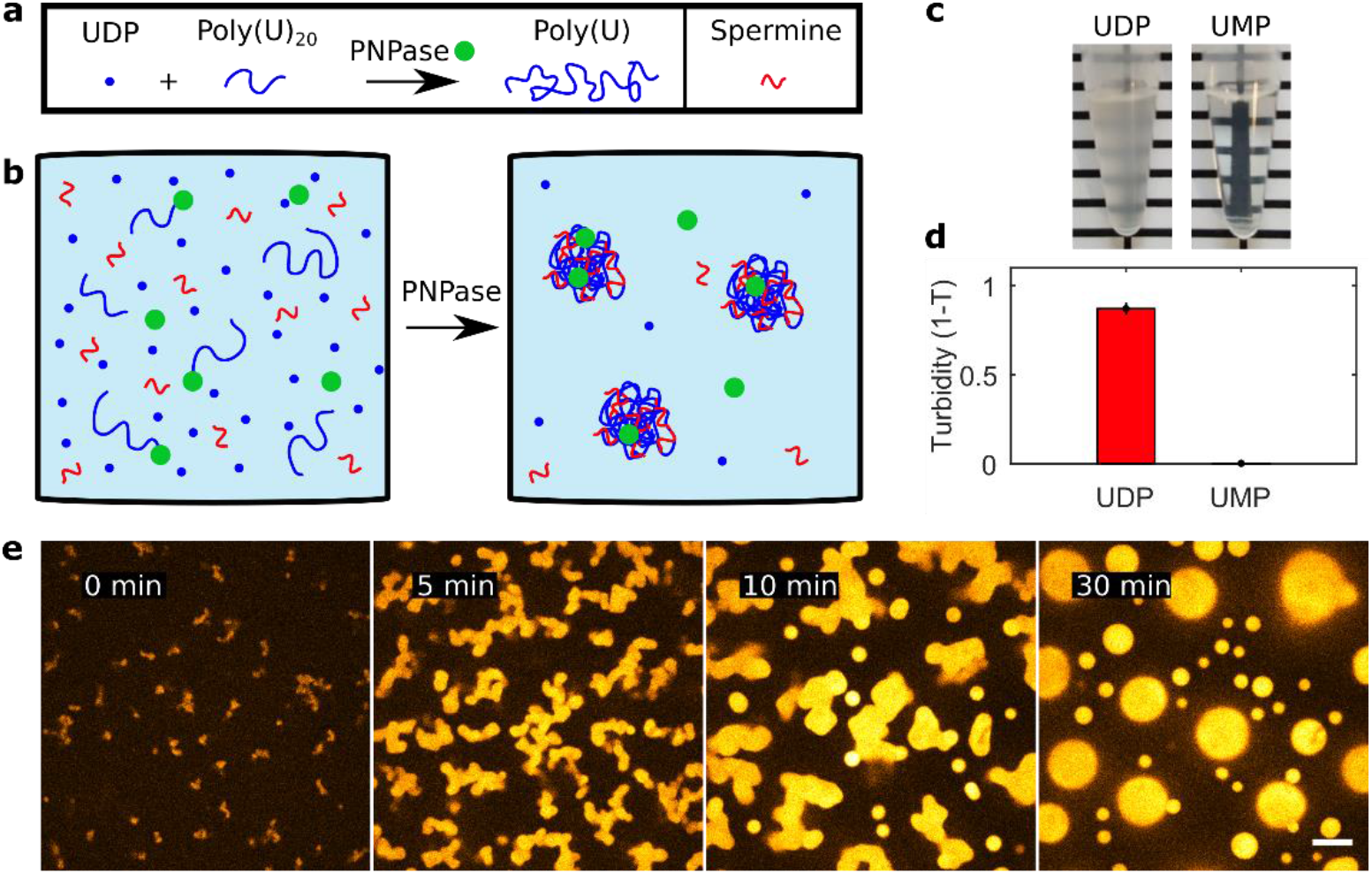
Poly(U)/spermine coacervates can be generated and activated by the enzyme Polynucleotide Phosphorylase (PNPase). (a) Schematic of PNPase RNA polymerization. PNPase (green circle) polymerizes short RNA seeds (blue lines) by adding UMP monomers from UDP (blue dots), giving rise to long homopolymeric polyuridylic RNA (poly(U)). (b) Schematic description of PNPase induced coacervation. Poly(U)_20_ does not phase separate within the presence of the polycation spermine, but once elongated, phase separation is initiated. (c) Coacervation induced by PNPase polymerization of poly(U) can be visualized macroscopically through solution turbidity. (d) Turbidity measurements of the solutions in panel c. Turbidity values were calculated from absorbance measurements at 500 nm wavelength (n = 9). (e) Time-lapse micrographs of the experiment shown schematically in panel b. The reaction contained 4 µM PNPase, 40 mM UDP, 1.0 wt% spermine and 5 µM Cy5-U20 and was carried out at 30 °C. Images are false-colored and the scale bar indicates 10 µm.

### Active coacervates initially assume non-spherical shapes

To better understand the observation of non-sphericality, we systematically tested various experimentally accessible parameters of the coacervate reaction. The parameters studied were nucleotide and enzyme concentrations, as well as temperature dependence. These parameters are expected to influence the reaction kinetics, the viscoelastic properties of the coacervate phase, and the chemical properties of the solution. All reactions contained 1.0% (w/v) spermine, 0.1% (w/v) poly(U), 5 µM Cy5-poly(U)_20_ seeds, and various amounts of UDP and PNPase (Fig. 2a). We expected low enzymatic activity at low UDP- and PNPase concentrations, corresponding to the cases of limiting substrate concentration (UDP) and when PNPase concentration was not sufficient to polymerize significant amounts of RNA in excess of 3’-ends for polymerization. The effect of temperature is more complicated since both the enzyme activity and the viscoelastic properties of the coacervates are likely affected. The coupling between these two properties is unknown. There were considerable qualitative differences between the shapes of coacervates activated under different conditions. For example, the coacervates activated by 1 µM PNPase appeared significantly more spherical than the ones activated by 4 µM PNPase (Fig. 2b & c). This qualitative difference in morphology is related to the interfacial energy associated with the coexistence of the two phases at hand, because the interfacial energy of an interface between two coexisting liquid phases is proportional to the interfacial surface area, with the interfacial tension being the constant of proportionality^43^. It is known that properties such as polymer length and the ionic strength of the solution affect the interfacial tension of coacervate-solvent interfaces^44,45^. In our case, these properties are dynamic, so that the interfacial tension is likely not a constant but actually varies over time in an unknown manner. Yet, the deviation from a spherical shape indicates that a non-minimal interfacial energy is associated with the coacervate-solvent interface. In an effort to quantify such differences between reaction conditions, we measured the average circularity of the coacervate cross sections. Circularity (𝜗) is a measure for how closely a two-dimensional object resembles a circle. It is defined through 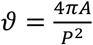, where *A* is the area and *P* the perimeter of the object. A circularity of 𝜗 = 1 implies that the object is a perfect circle, whereas a circularity of 𝜗 = 0 implies an infinitely long and infinitely thin line. In three dimensions, the circularity of a single cross section cannot be related one-to-one to the total energy of the surface. However, we argue that the average circularity of cross sections obtained for multiple coacervates provides an indirect measure of how far the coacervates are from their spherical equilibrium shape (see **Supporting Information**).

**Figure 2:**
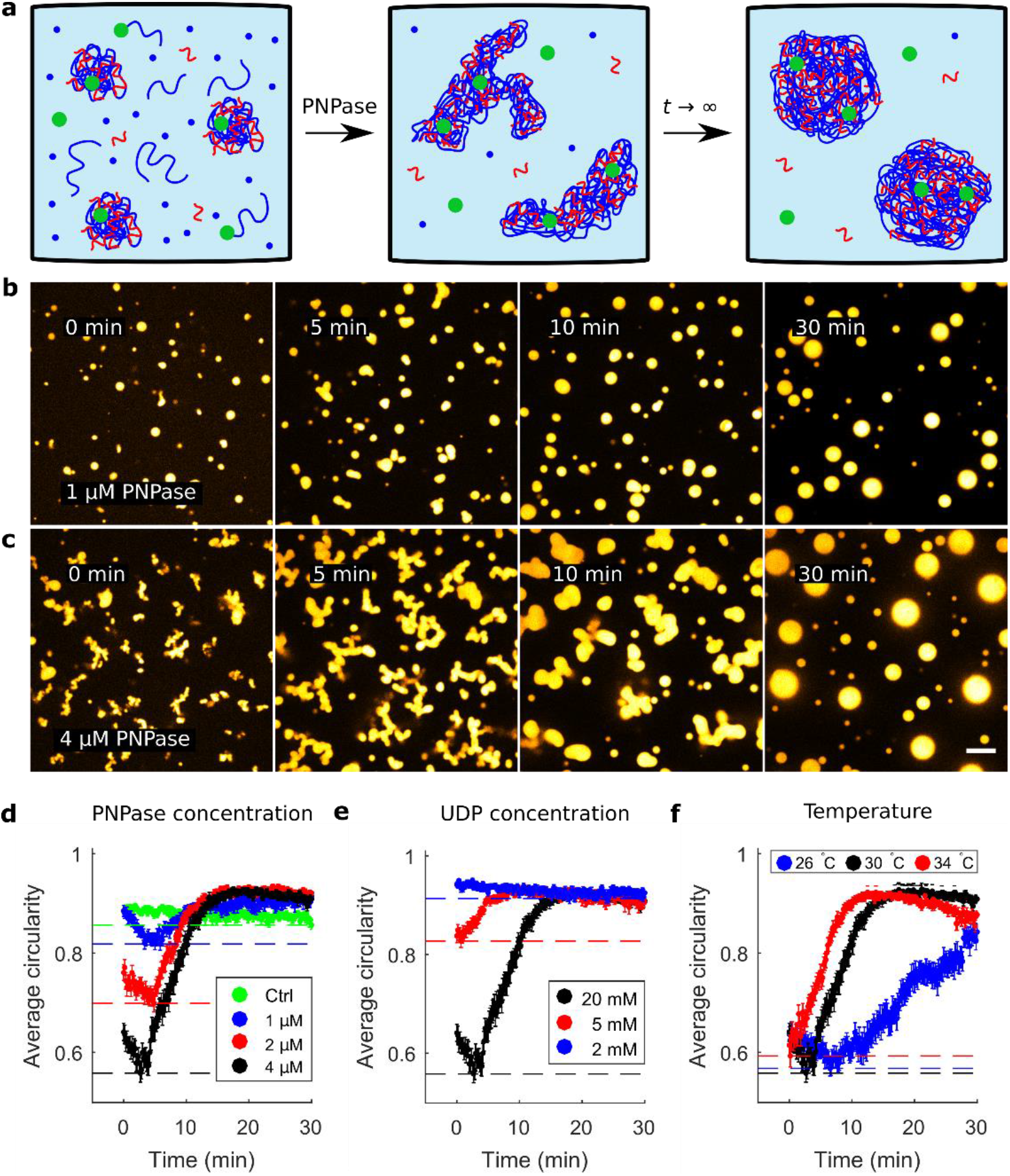
Quantification of non-equilibrium coacervate shapes by average circularity of cross sections. (a) Schematic depiction of PNPase reaction in the presence of poly(U)/spermine coacervates in equilibrium, supplemented with poly(U) seeds, UDP and PNPase. Initially, the coacervates become active but as the reaction reaches a dynamic equilibrium (*t* → ∞), the coacervates assume their spherical equilibrium shape. (b, c) PNPase reaction in the presence of pre-formed coacervates with 1 µM (b) and 4 µM (c) PNPase. There is a significant qualitative difference in terms of the coacervate shapes prior to assuming a spherical equilibrium shape. Images are false-colored and the scale bar indicates 10 µm. (d) Plot of the average circularity of coacervate cross sections for PNPase concentrations of 4 µM (black), 2 µM (red) and 1 µM (blue) PNPase. The control (green) contained passive poly(U)_∞_/spermine coacervates, which was produced by incubation of PNPase, poly(U)_20_ and UDP (see **Materials & Methods** for details). Dashed lines indicate minimum observed average circularity per condition. In all images, PNPase was added 5 min before the start of image acquisition. (e) Average circularity profiles for 20 mM (black), 5 mM (red) and 2 mM (blue) UDP. (f) Average circularity profiles for temperatures of 30 °C (black), 34 °C (red) and 26 °C (blue). In panels d-f the error bars indicate standard error from the mean.

To assess PNPase induced non-sphericality of coacervates under different conditions, the micrograph images of PNPase-activated coacervates were binarized (**Materials & Methods**), and the average circularity was determined. The average circularities are plotted over time in Figure 2d-f for the various PNPase concentrations, UDP concentrations and temperatures. Qualitatively, the circularity profile for active coacervates was similar across all conditions. Initially, the average circularity decreases to a Minimum Average Circularity (the MAC-value). This MAC-value is indicative of how much the coacervates deviate from their spherical equilibrium shape. After reaching the MAC-value, the average circularity recovers to a constant value (𝜗 > 0.85). We noted that a higher PNPase concentration correlated with a lower MAC-value, implying that higher PNPase concentrations take coacervates further away from their equilibrium shapes. However, the time in which the coacervates regained spherical shapes was comparable for all tested PNPase concentrations (Fig. 2d). Since an increase in PNPase concentration does not affect the relaxation time into spherical coacervates, it appears that physical properties of the system are the main determinants of the relaxation process. Similarly, we tested the effect of UDP concentration. A UDP concentration of 20 mM yielded a significantly lower MAC-value than concentrations of 5 and 2 mM (Fig. 2e). Therefore we establish that at higher UDP concentrations, the coacervates are initially taken further out of their spherical equilibrium shape.

In contrast to variations in PNPase concentration, the time in which the coacervates become spherical does differ between UDP concentrations, where non-spherical shapes persist for longer times at higher UDP concentration. Note that higher UDP (disodium salt) concentrations may also have the opposite effect, because beyond a critical salt concentration coacervates dissolve^36^. Taken together, the titration experiments of PNPase and UDP suggest that the degree to which active coacervates are non-spherical (as indicated by the MAC-value) is correlated to the enzyme activity.

Finally, we assessed the effect of temperature on the shapes of active coacervates. We observed only small differences at temperatures of 30 and 34 °C in the average circularity profile. At 26 °C, the average circularity reached a MAC-value comparable to that observed at the other temperatures, but it took significantly longer to recover to a steady state value (Fig. 2f). This difference may be explained by the fact that the phase behavior of the system strongly depends on temperature. Aumiller Jr. and colleagues have demonstrated that the minimum temperature for RNA/spermine coacervation is around ~27 °C^36^.

To conclude this section, we have observed a clear correlation between the degree to which coacervates were driven away from their spherical equilibrium shape (as measured by the MAC-value), PNPase concentration, and UDP concentration. Our observations are in agreement with the assumption that increasing PNPase and UDP concentrations increases the net poly(U) polymerization rate. Therefore, we conclude that we have found a positive correlation between overall poly(U) polymerization rate and the degree to which the activated coacervates are taken out of equilibrium. For the temperatures we tested, the MAC-values were similar but the relaxation to spherical shape happened substantially slower at 26 °C. This means that temperature affects the PNPase activity and the viscoelastic properties of the coacervate phase, with an unknown coupling, which precludes a simple explanation for our observation.

### Non-spherical coacervates persist on a timescale significantly exceeding that of coacervate merging

To better understand the phenomenon of non-sphericality we asked the question of whether the observed coacervate shapes are related to viscoelastic properties of the coacervate phase. One way to determine viscoelastic properties of liquid phases is to quantify the relation between the timescale of droplet merging events, and droplet size^46,47^. To first order, the timescale of convective flow (*T*) driven by interfacial tension between liquids, is proportional to the radius of coacervate droplets (*L*). The inverse capillary velocity is the proportionality constant for the relation *T* ∝ *L*, and is defined as the ratio between the viscosity *η* and the interfacial tension *γ*, which are both material properties that also depend on temperature. To determine the timescale of merging at given length scales, we developed an image processing algorithm which quantifies the Circularity Recovery After Merging (CRAM). CRAM-analysis consists of the acquisition of coacervate merging events and examining the circularity of the resulting coacervate over time (Fig. 3a). The recovery of circularity after merging of two coacervates follows approximately an exponential function. By fitting an exponential function to the circularity of individual merging events, we quantified the timescale of merging (Fig. 3b and **Materials & Methods**). We considered merging events of spherical coacervates of approximately equal size that are smaller than 4 µm in diameter. The correlation between time- and length scales (coacervate radius) of the merged coacervate, yielded an inverse capillary velocity of 0.67 s/µm (R^2^ = 0.53, n = 53, Fig. 3c). This value is two orders of magnitude lower than, for example, the inverse capillary velocity of *Xenopus laevis* nucleoli, which was found to be 46.1 s/µm^46^. Importantly, the timescale resulting from the inverse capillary velocity is two orders of magnitudes faster than the timescale at which non-spherical PNPase-activated coacervates persist (~10 min, Figs. 1e, 2d-f). The timescale analysis allows us to conclude that non-sphericality cannot be explained by merging spherical coacervates but arises from enzymatic activity of PNPase and its polymerization of poly(U) RNA. This view is additionally supported by the mobility of RNA polymers inside coacervates. Figure 3d shows a merging event of two (passive) coacervates that had been prepared with two different fluorescent labels on the poly(U) seeds, Cy5 and Cy3. We observed that RNA inside the coacervate phase is mobile and rapidly mixes within seconds, in support of rapid diffusive mixing of poly(U) inside the coacervates (**Fig. S7**).

**Figure 3:**
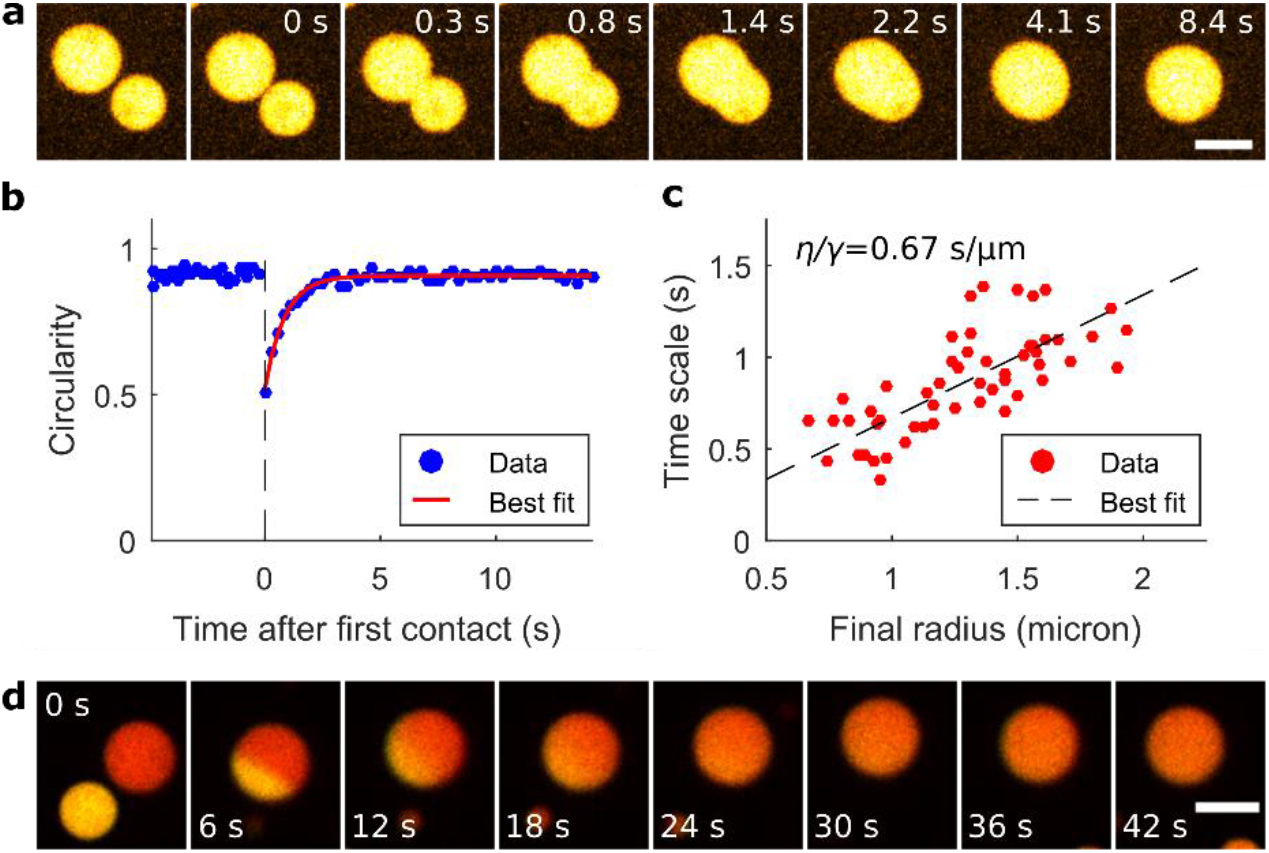
Analysis of the Circularity Recovery After Merging (CRAM) provides the timescale of coacervate merging. (a) Micrograph images of a merging event. The first image shows two coacervates prior to the merging process. Time zero is defined as the moment when the coacervates start merging. (b) Circularity-profile corresponding to the merging event of panel a. Prior to merging (*t* < 0), two spherical coacervates are observed. As they merge, the circularity of the resulting coacervate drops sharply, and recovers to a stationary value as the coacervates regains spherical shape. The red line indicates the best fit of a single exponential (see **Materials & Methods**) (c) Scatterplot of the timescale of merging plotted against the radius of the resulting droplet. The inverse capillary velocity was found to be 0.67 s/µm by linear regression (R^2^ = 0.53, n = 53), indicated by the black dashed line. (d) Merging of passive coacervates containing Cy3- and Cy5-poly(U) (yellow and red respectively, images false-colored) to demonstrate the mobility of the poly(U) within the coacervate phase. Scale bars of panels a and d indicate 5 µm.

### PNPase localizes predominantly inside the coacervate phase

To determine the localization of PNPase with respect to the coacervate phase and the aqueous phase, we purified and chemically labeled PNPase with FITC (**Supporting Information**). Using confocal microscopy FITC-PNPase was observed to form aggregates which localize to the coacervate-solvent interface (**Fig. S5**). Sometimes multiple coacervates were connected to each other through PNPase patches. To accurately assess the partitioning of fully solubilized FITC-PNPase in the poly(U)/spermine coacervate system, we used ultracentrifugation (Airfuge, 5 min, 100,000 g) of the FITC-PNPase stock prior to adding FITC-PNPase to the sample. We observed a significant decrease in the amount of FITC-PNPase aggregates before the addition of UDP. We observed that PNPase localizes predominantly in the coacervate phase, but that there is significant fluorescence intensity from the surrounding solvent phase (Fig. 4a & b). We calculated the fluorescence intensity inside and outside the coacervate phase by using the Cy5-poly(U) fluorescence intensities as a mask to define the phases (see **Materials & Methods** and **Fig. S6**). The FITC-PNPase concentration was approximately 3.9-fold higher inside the coacervate phase compared to the solvent phase (Fig. 4c). We also tested if FITC-PNPase had a preference for the interface between the active coacervate and the solvent, as suggested by the observation of aggregates sticking to the coacervate surface. However we did not find significant accumulation of PNPase at the edge of the coacervate phase (Fig. 4c). Besides enzyme localization, we also tested if the coacervate phase was permeable to FITC-PNPase. To this end we performed Fluorescence Recovery After Photobleaching experiments (FRAP, see **Materials & Methods**), and observed that a fully bleached coacervate recovers FITC-PNPase fluorescence within several minutes. It can be concluded that there is continuous exchange of FITC-PNPase between the coacervate and solvent phase, and that there is rapid internal redistribution of FITC-PNPase throughout the coacervate phase.

**Figure 4:**
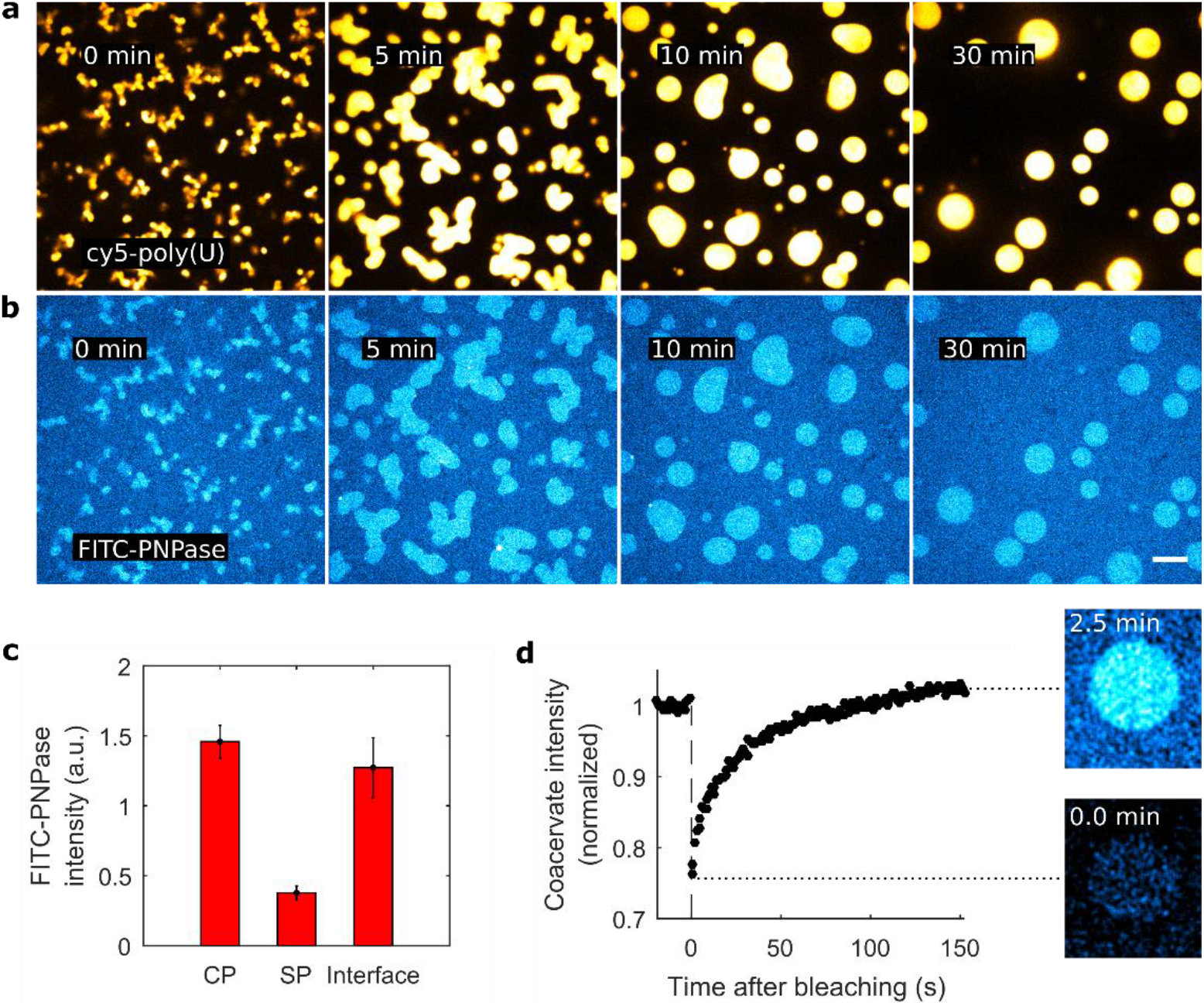
Localization of reaction components in active poly(U)/spermine coacervates. (a, b) PNPase reaction with 5’-end Cy5-labeled poly(U) (panel a) and FITC-labeled PNPase (panel b). To illustrate the difference, images were false colored with Cy5-poly(U) in yellow and FITC-PNPase in blue. Scale bar indicates 10 µm. (c) Fluorescence intensities of FITC-PNPase in the coacervate phase (CP), solvent phase (SP) and at the CP-SP interface. This implies that the PNPase concentration inside the coacervate phase is 3.9-fold higher than in the surrounding solvent phase (see **Materials & Methods**). Error bars indicate the standard deviation. (d) Fluorescence Recovery After Photobleaching (FRAP) of FITC-PNPase fluorescence indicates recovery of FITC-PNPase within 2.5 minutes. This indicates that there is continuous exchange of PNPase with the surrounding solvent phase and that the PNPase is redistributed throughout the coacervate phase.

### Coacervate degradation by PNPase in high-phosphate buffer

Besides poly(U) polymerization by incorporating UMP monomers from free UDP, PNPase also catalyzes the reverse reaction (Fig. 5a). We attempted to degrade poly(U)/spermine coacervates by supplying excess phosphate and PNPase (Fig. 5b), and indeed observed significant degradation of coacervates within 30 minutes (Fig. 5c). This shows that poly(U)/spermine coacervates can both be generated, as well as degraded depending on the phosphate levels of the buffer. Thus, we hypothesize that the PNPase reaction reaches a steady state of spherical coacervates in which polymerization and depolymerization reactions have reached a dynamic chemical equilibrium.

**Figure 5:**
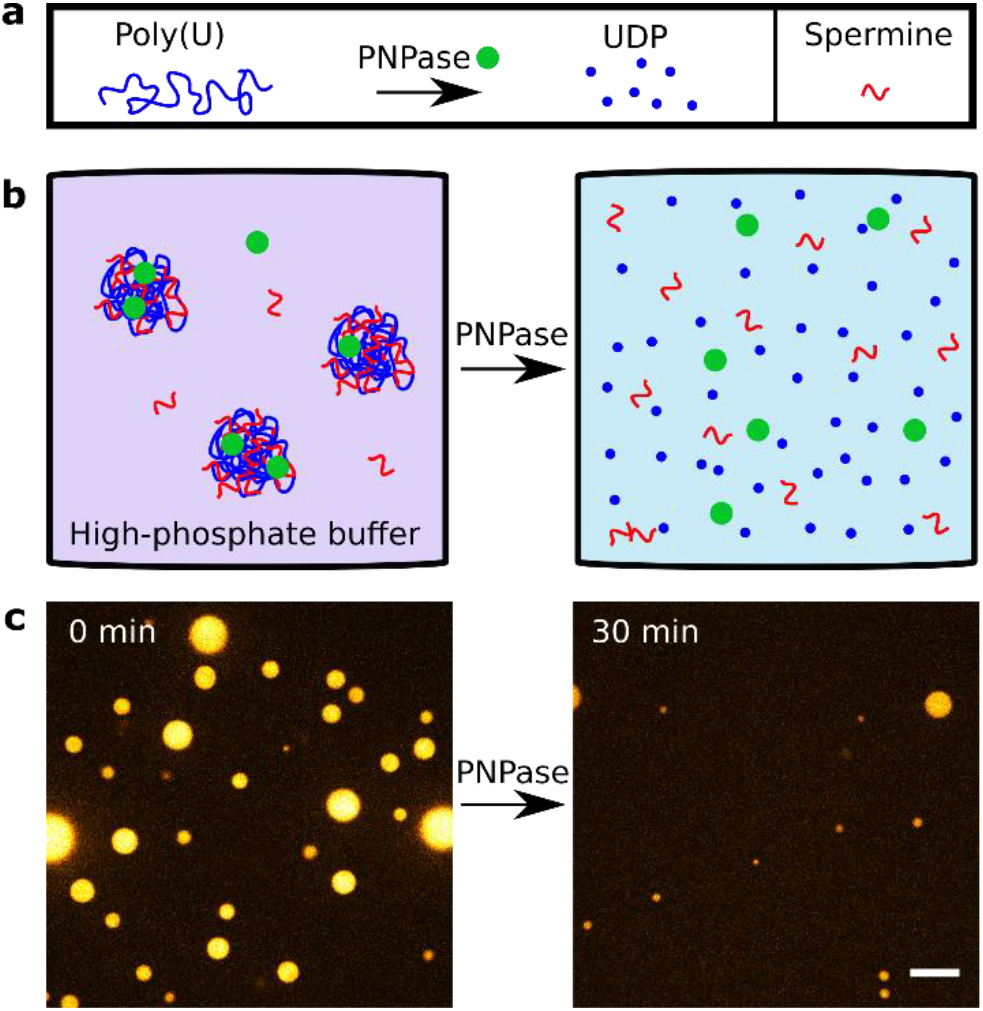
PNPase degrades coacervates in high-phosphate buffer. (a) Provided with phosphate instead of nucleotides, PNPase has 3’-to-5’ exoribonuclease activity, through which it forms UDP from poly(U) and free phosphate. (b) Implementation of PNPase-degradation of coacervates inside high-phosphate buffer. (c) Micrograph images of the experiment shown in panel b. Poly(U)_∞_/spermine coacervates were prepared in a buffer containing 10 mM disodium phosphate and 4 µM PNPase. After 30 minutes significant degradation of coacervate was observed. Images are false-colored and the scale bar indicates 10 µm.

## Discussion

Inspired by the observation that biological systems are highly dynamic and generically active, we have presented and characterized an example of an active RNA/spermine coacervate system. The activity of the coacervates was provided by the enzyme PNPase, which polymerizes RNA templates. Unexpectedly, we observed that the shapes of coacervates initially deviated from the spherical equilibrium shape. Because non-spherical coacervates are a phenomenon emerging from combining the RNA polymerization and its coacervation with polycations, it is not possible to attribute these morphological changes to either process as the leading cause.

Our findings may help to establish experimental model systems and to develop and test theories of chemically/enzymatically active coacervation. To this end, a set of experiments has been conducted to quantify and characterize the system’s phenomenology. We have found that the extent to which coacervate shapes deviate from spherical shape (as measured by the MAC-value) correlates with higher enzyme activity. The timescale at which active coacervate shapes are non-spherical is two orders of magnitude longer than the timescale at which two passive coacervates merge to one spherical coacervate. While the spherical coacervate shape is quickly recovered after merging, this does not necessarily imply that the coacervate material mixes well. As time progresses, longer RNA molecules may prevent mixing of coacervate material, despite quick merging events. This can be attributed to a reduced diffusion coefficient of very long RNA molecules inside the coacervate phase. Notably, also the chemical environment in the active coacervate system changes over time and influences its physical properties.

Overall, PNPase activity affects the viscoelastic properties of the coacervate phase in a variety of ways. Firstly, the RNA is being elongated and it is established that the length of polymers is an important factor in its ability to coacervate. Additionally, this elongation decreases the RNA diffusivity both inside the coacervate phase, as well as in the dilute phase. Secondly, the spermine/poly(U) ratio changes dynamically and significantly affects the physico-chemical properties of coacervates in the course of time. Thirdly, there is a rising level of phosphate from the incorporation of UMP monomers from UDP. This affects the ionic strength and the pH of the solution, and could be quantified in future experiments (e.g. using nuclear magnetic resonance spectroscopy or thin layer chromatography). In addition, we note that chemical gradients may arise inside the system, for example due to differential localization of PNPase between the phases. It is unclear how these gradients affects the viscoelastic properties of the coacervates. Accurate quantification of the viscoelastic properties of the coacervates will be useful in determining the precise physical state of the observed structures, which ranges from liquid to more gel-like and solid. One may speculate that the high initial ratio of spermine to poly(U) causes the coacervates to have more gel-like properties during the formation of structures. The PNPase reaction causes the spermine/poly(U) ratio to decrease due to poly(U) polymerization. This decrease may cause the coacervates to become more liquid and even dissolve when the charge ratio drops below a critical point^36^.

We would like to highlight two follow-up approaches to further elaborate our findings. First, it would be interesting to look at the underlying physical mechanisms that lead to non-spherical shapes. There exist several theoretical models for the dynamics of LLPS, such as the stochastic and/or advective Cahn-Hilliard-Cook models^13^, which describe binary de-mixing processes. Furthermore, theory allows the investigation of complex reaction-diffusion systems in which chemical reactions drive shape changes of active coacervates^31,32^. Although some of these theoretical models predict the formation of non-spherical coacervates at the initial stages of liquid-liquid phase separation, most verified models lack the explicit incorporation of (enzymatic) activity and how this affects the physicochemical properties of the reaction components during the process^13^. Specifically, the system we studied is highly complex because it depends on a variety of dynamic quantities mentioned above. The incorporation of factors such as enzyme activity, polymer length, ionic strength, and charge ratio into existing theoretical models may reveal potential mechanisms underlying our observations. Understanding the mechanisms based on physical and chemical processes will be an interesting avenue for further research.

A second approach would be to more closely investigate the non-equilibrium states of the system and determine the coupling between the processes of enzymatic RNA polymerization and coacervation^48^. While each of the two processes on their own reach thermodynamic equilibrium directly (i.e. there is a single minimum in the Gibbs free energy), the combined system displays an interesting deviation from the classical picture. The system needs constant input of energy to maintain its dissipative non-equilibrium state. As the chemical composition of the system changes it may transition through a variety of non-equilibrium states before reaching thermodynamic equilibrium.

## Conclusion

As a final note we wish to mention that our study may provide a simple model system for a variety of biological and chemical processes. Besides serving as a model for protocells, as originally suggested by Oparin^15–17^, we see at least two more applications towards biological systems. First of all, membrane-less organelles have been shown to be (active) biomolecular condensates^8^. Some intracellular biomolecular condensates have been shown to dynamically display non-equilibrium shapes. One notable example is the pyrenoid in *Chlamydomonas reinhardtii*, which is involved in carbon fixation during photosynthesis^49^. Another application of active coacervates is the bottom-up engineering of synthetic lifeforms^50–53^, which also requires compartmentalization and chemical activity as provided in parts by our system. Such efforts frequently include combinations of both membranous (e.g. liposomes) and membraneless (e.g. coacervates)^54–57^ compartments. In order to better understand biological systems, such as (parts of) prebiotic protocells, contemporary cells or future synthetic lifeforms, it will be necessary to study the characteristics and uncover fundamental laws that govern active liquid-liquid phase separation.

## Acknowledgements

Plasmid pET20b, containing C-terminally His_6_-tagged PNPase was kindly provided by Gadi Schuster, Technion University, Faculty of Biology (Israel). We thank Siddharth Deshpande, Anson Lau and Cees Dekker for fruitful discussions. We furthermore thank Elena S. Nadezhdina, Institute of Protein Research (Russian Academy of Science), for providing Oparin’s original papers in Russian, and Vladimir Volkov for his help with translating. L. Reese was supported by FOM programme nr. 110 (NWO). E. O. van der Sluis was supported by Sinergia grant 160728 from the Swiss National Science Foundation (SNF).

## Author contributions

W.K.S., L.R. and M. D. conceived the study. W.K.S. performed the experiments and analyzed the data. E.O.v.d.S. purified and labelled the PNPase. All authors discussed the data. L.R. and M.D. supervised the research. W.K.S. and L.R. wrote the manuscript with input from all authors.

## Supporting Information

Supporting Information is available containing additional text, figures and videos.

## Table of Contents Graphic

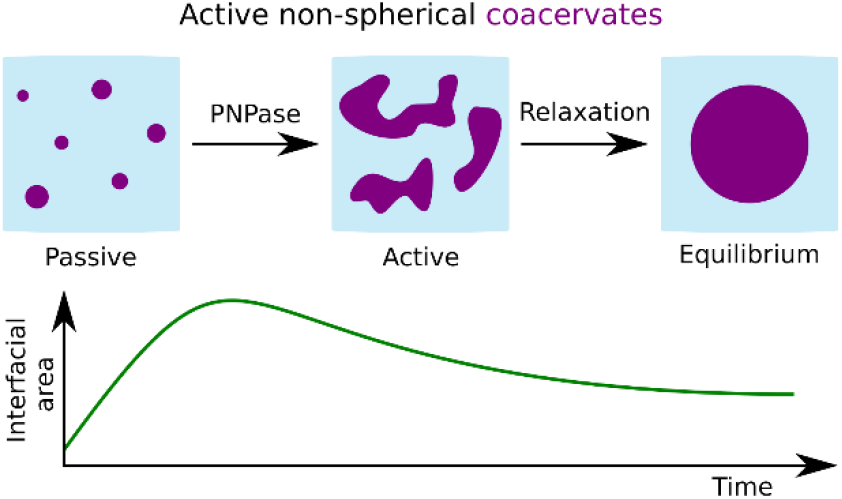

